# Leveraging TOPMed Imputation Server and Constructing a Cohort-Specific Imputation Reference Panel to Enhance Genotype Imputation among Cystic Fibrosis Patients

**DOI:** 10.1101/2021.12.20.473535

**Authors:** Quan Sun, Weifang Liu, Jonathan D. Rosen, Le Huang, Rhonda G. Pace, Hong Dang, Paul J. Gallins, Elizabeth E. Blue, Hua Ling, Harriet Corvol, Lisa J. Strug, Michael J. Bamshad, Ronald L. Gibson, Elizabeth W. Pugh, Scott M. Blackman, Garry R. Cutting, Wanda K. O’Neal, Yi-Hui Zhou, Fred A. Wright, Michael R. Knowles, Jia Wen, Yun Li, on behalf of the Cystic Fibrosis Genome Project

## Abstract

Cystic fibrosis (CF) is a severe genetic disorder that can cause multiple comorbidities affecting the lungs, the pancreas, the luminal digestive system and beyond. In our previous genome-wide association studies (GWAS), we genotyped ∼8,000 CF samples using a mixture of different genotyping platforms. More recently, the *Cystic Fibrosis Genome Project* (CFGP) performed deep (∼30x) whole genome sequencing (WGS) of 5,095 samples to better understand the genetic mechanisms underlying clinical heterogeneity among CF patients. For mixtures of GWAS array and WGS data, genotype imputation has proven effective in increasing effective sample size. Therefore, we first performed imputation for the ∼8,000 CF samples with GWAS array genotype using the TOPMed freeze 8 reference panel. Our results demonstrate that TOPMed can provide high-quality imputation for CF patients, boosting genomic coverage from ∼0.3 - 4.2 million genotyped markers to ∼11 - 43 million well-imputed markers, and significantly improving Polygenic Risk Score (PRS) prediction accuracy. Furthermore, we built a CF-specific *CFGP reference panel* based on WGS data of CF patients. We demonstrate that despite having ∼3% the sample size of TOPMed, our *CFGP reference panel* can still outperform TOPMed when imputing some CF disease-causing variants, likely due to allele and haplotype differences between CF patients and general populations. We anticipate our imputed data for 4,656 samples without WGS data will benefit our subsequent genetic association studies, and the CFGP reference panel built from CF WGS samples will benefit other investigators studying CF.

## Introduction

Cystic fibrosis (CF) is an autosomal recessive genetic disorder caused by mutations in the *cystic fibrosis transmembrane conductance regulatory* (*CFTR*) gene. CF affects the lungs, pancreas, and other organs, but the major cause of morbidity and mortality is progressive obstructive lung disease and lung injury due to inflammation and infection. We previously have conducted genome-wide association studies (GWAS) for CF and related traits^1–4^, where we genotyped ∼8,000 CF samples at approximately half a million common genetic variants, imputed up to 8.5 million markers using haplotypes combined from the 1000 Genomes Project and deep (∼30X) sequence from 101 Canadian CF patients as reference, and evaluated association between each genotyped or imputed marker and CF or related traits.

Recently, our *Cystic Fibrosis Genome Project* (CFGP) generated high-coverage (∼30X) whole genome sequence (WGS) data for 5,095 CF samples. Together with our previous GWAS efforts, we have 1,880 CF samples with WGS data alone, 4,656 samples with GWAS data alone, and 3,215 patients with both WGS (3,215 samples) and GWAS data (3,314 samples, due to sample duplicates/triplicates). In this work, we set out to ask two questions. First, would the latest imputation reference panel from the NHLBI Trans-Omics for Precision Medicine (TOPMed) project aid imputation among CF patients? TOPMed has demonstrated its value in further boosting imputation quality and rescuing lower frequency and rare variants due to its large sample size representing diverse ancestries^5,6^. We hypothesize that CF patients may similarly benefit from the TOPMed imputation reference panel. Second, is there any value in building a CF-specific reference panel based on WGS data from CF patients? For example, the CF-causing 3bp deletion c.1521_1523delCTT [p.Phe508del; legacy name: F508del] in *CFTR* has a frequency of 69.7% among CF patients (CFTR2) but merely 0.8-1.0% in general populations across continental groups (Bravo). We hypothesize that a CF-specific reference panel may better recover CF associated regions, even though the TOPMed sample size (n=97,256) is ∼20X that in CFGP (n=5,095), given the presumably more drastic allele and haplotype pattern differences at CF related loci. For the second question, Panjwani et al^7^ showed the value of including CF patients in imputation reference panel, where they included haplotypes from a much smaller set (n=101) of CF patients. Systematic comparisons with larger sample sizes are still lacking.

In this manuscript, we first performed imputation of different CF datasets starting from array genotype only, leveraging the TOPMed freeze 8 reference panel. We then systematically evaluated the imputed data using the WGS data as the working truth. Evaluations included quantifying the number of well-imputed variants, assessing the true imputation quality, gauging heterozygous concordance for extremely rare variants, and evaluating imputation quality for the *CFTR* F508del variant in comparison with previous work^7^. We then constructed a *reduced-CFGP* reference panel to evaluate if the WGS data of CF patients would provide additional insights beyond TOPMed-based imputation. Finally, we constructed PRS for KNoRMA, a lung function measurement, to assess the impact of imputation on PRS construction.

In this paper, we refer to observed genotypes derived from WGS data as “true genotypes”, though in reality genotype calls from WGS data are not 100% accurate. We use “true R2” (**Method**) to refer to the squared Pearson correlation between imputed dosages and “true genotypes” from WGS data, and use “Rsq” output from imputation software to denote the estimated imputation quality. Note that the calculation of “true R2” entails “true genotypes” which we do not have in typical imputation while Rsq is available whenever imputation is performed.

## Results

### Imputation with TOPMed freeze 8 reference panel and quality evaluation

To answer how the TOPMed reference panel would aid imputation in CF, we imputed 7,970 CF samples with genotyping array data, leveraging the imputation reference panel built from 97,256 deeply-sequenced human genomes in the TOPMed project. These 7,970 samples were genotyped using various commercial genotyping platforms directly examining 263,660 - 4,389,087 variants, in various projects including the CF Twin and Sibling Study, the CF-related Diabetes (CFRD) Study, the Gene Modifier Study (GMS), and the GMS CF Liver Disease Study^1–4^. For a subset of 2,933 samples with WGS data from the CFGP, we then assessed the imputation quality by comparing imputed dosages to observed genotypes in the WGS data, with the latter treated as the gold-standard.

We focused on two metrics in our imputation quality evaluation: the number of well-imputed variants and average imputation quality for these well-imputed variants. We first assessed the numbers of well-imputed variants by minor allele frequency (MAF) separately for the seven GWAS arrays. We applied post-imputation quality filtering, based on estimate R^2^ (or Rsq), using two different thresholds (Rsq >= 0.3 or Rsq >= 0.8 with the latter being the more stringent/aggressive filtering). Both thresholds are commonly adopted for post-imputation quality filtering^8–10^. Using the TOPMed reference panel, we obtained 11,156,390 - 43,095,581 well-imputed variants (Rsq >= 0.8) including 2,533,058 - 33,399,492 low frequency or rare variants (LFRV; MAF <= 0.5%) (**Table 1**). For example, for the 3,840 samples genotyped with the Illumina 610-Quad array, we observed 43,095,581 well-imputed (Rsq >= 0.8) variants with 33,399,492 being LFRV.

**Table 1.** Numbers of well-imputed variants by different MAF categories for the seven GWAS arrays (genome wide)

| Illumina Panel<sup>a</sup> | Number of samples<sup>a</sup> | Number of samples-by-site<sup>a</sup> | Number (%)<sup>b</sup> of SNPs Rsq≥ 0.3 | Number (%)<sup>b</sup> of SNPs Rsq ≥0.8 | Number (%)<sup>c</sup> of SNPs Rsq ≥0.8 & MAF<0.5% | Number (%)<sup>d</sup> of SNPs Rsq ≥0.8 & MAF<5% | Number (%)<sup>e</sup> of SNPs Rsq≥ 0.8 & MAF>=5% |
| --- | --- | --- | --- | --- | --- | --- | --- |
| 300K | 144 | FrGMC<br>1,300 | 17,603,215<br>(5.73%) | 12,248,616<br>(3.99%) | 3,897,584<br>(1.31%) | 6,738,025<br>(2.24%) | 5,510,591<br>(88.02%) |
| 370K | 145 |  | 14,471,514<br>(4.71%) | 11,156,390<br>(3.63%) | 2,533,058<br>(0.85%) | 5,519,937<br>(1.83%) | 5,636,453<br>(90.49%) |
| 660K | 1,011 |  | 30,661,930<br>(9.99%) | 20,830,921<br>(6.79%) | 11,883,847<br>(4.01%) | 15,138,988<br>(5.03%) | 5,691,933<br>(93.95%) |
| 610-Quad | 3,840 | CGS 1,533;<br>GMS 1467;<br>TSS 840 | 58,672,809<br>(19.12%) | 43,095,581<br>(14.04%) | 33,399,492<br>(11.26%) | 37,276,108<br>(12.39%) | 5,819,473<br>(96.22%) |
| 660W-set1 | 2,012 | CGS 342;<br>GMS 808;<br>TSS 862; | 43,832,169<br>(14.28%) | 34,503,481<br>(11.24%) | 24,694,173<br>(8.33%) | 28,669,926<br>(9.53%) | 5,833,555<br>(96.33%) |
| 660W-set2 | 444 | TSS 444 | 23,814,328<br>(7.76%) | 20,792,798<br>(6.77%) | 10,176,358<br>(3.43%) | 14,916,691<br>(4.96%) | 5,876,107<br>(96.98%) |
| Omni5 | 374 | CGS 73;<br>GMS 170<br>TSS 131; | 20,774,826<br>(6.83%) | 18,862,492<br>(6.20%) | 10,530,015<br>(3.55%) | 14,053,383<br>(4.68%) | 4,809,109<br>(97.65%) |
| <sup>a</sup> Corvol et al 2015 reference<br><sup>b</sup> Percentage taken over total number of imputed variants from TOPMed freeze 8 reference panel<br><sup>c</sup> Percentage taken over imputed variants with MAF < 0.5%<br><sup>d</sup> Percentage taken over imputed variants with MAF < 5%<br><sup>e</sup> Percentage taken over imputed variants with MAF >= 5% |  |  |  |  |  |  |  |

We then calculated the average imputation quality for these well-imputed variants. Specifically, we calculated true R^2^ by comparing imputed dosages with WGS data which again serves as the “gold standard” (**Methods**). We evaluated two GWAS arrays with the largest sample sizes, Illumina 610-Quad and 660W-set1, to obtain a more stable imputation quality estimate for LFRV, and took chromosome 20 as an example. For samples genotyped with the 610-Quad array and 660W-set1, 1,992 and 941, respectively, also had WGS performed in the CFGP. Based on these 1,992 and 941 samples, we observed that average true R^2^ values for variants across all MAF categories are greater than 0.93, indicating that imputed dosages recover >93% information in the true genotypes (**Table 2**).

**Table 2.**
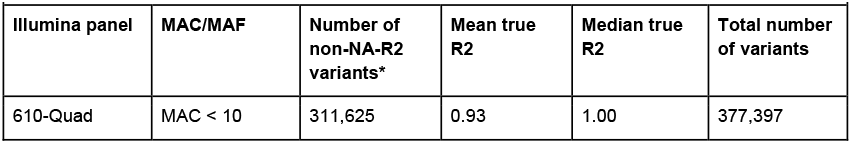
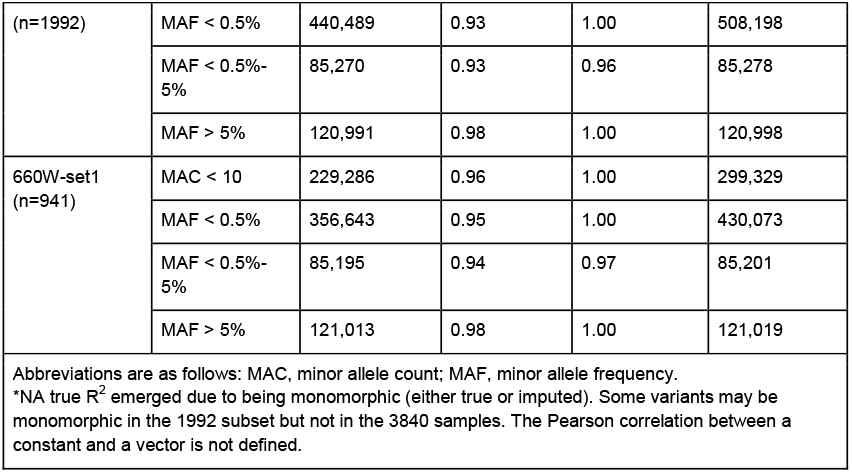
True R^2^ for the two arrays with the largest sample sizes (chr20)

We also gauged heterozygous concordance for extremely rare variants (defined as minor allele count, MAC, <10). Even for those extremely rare variants, the average heterozygous concordances are greater than 0.97 (**Table 3**), indicating that the TOPMed reference panel can impute those rare variants well. We specifically checked imputation quality for the *CFTR* F508del variant on chromosome 7 that, as aforementioned, has a drastic allele frequency difference between CF patients (69.7%) and general populations (0.8%). The estimated R^2^’s for 610-Quad and 660W-set1 arrays are 0.89 and 0.93 respectively; and the true R^2^’s are 0.83 and 0.87, suggesting that the imputation quality for this variant is rather decent, rescuing 83% and 87% of the information content. However, TOPMed reference panel tends to call the homozygote deletion genotype (1/1) as heterozygotes (0/1) (**Figure 1**), showing there is still room for improvement.

**Table 3.** Heterozygous concordance for extremely rare variants (chr20)

| **Table 3. Heterozygous concordance for extremely rare variants (chr20)** |  |  |  |  |  |
| --- | --- | --- | --- | --- | --- |
| **Illumina panel** | **Number of samples** | **Number of non-NA het concordant variants** | **Mean het concordant (freq)** | **Median het concordant (freq)** | **Total number of variants** |
| 610-Quad | 1992 | 212,759 | 0.98 | 1.00 | 296,088 |
| 660W-set1 | 941 | 289,811 | 0.97 | 1.00 | 374,166 |

**Figure 1.**
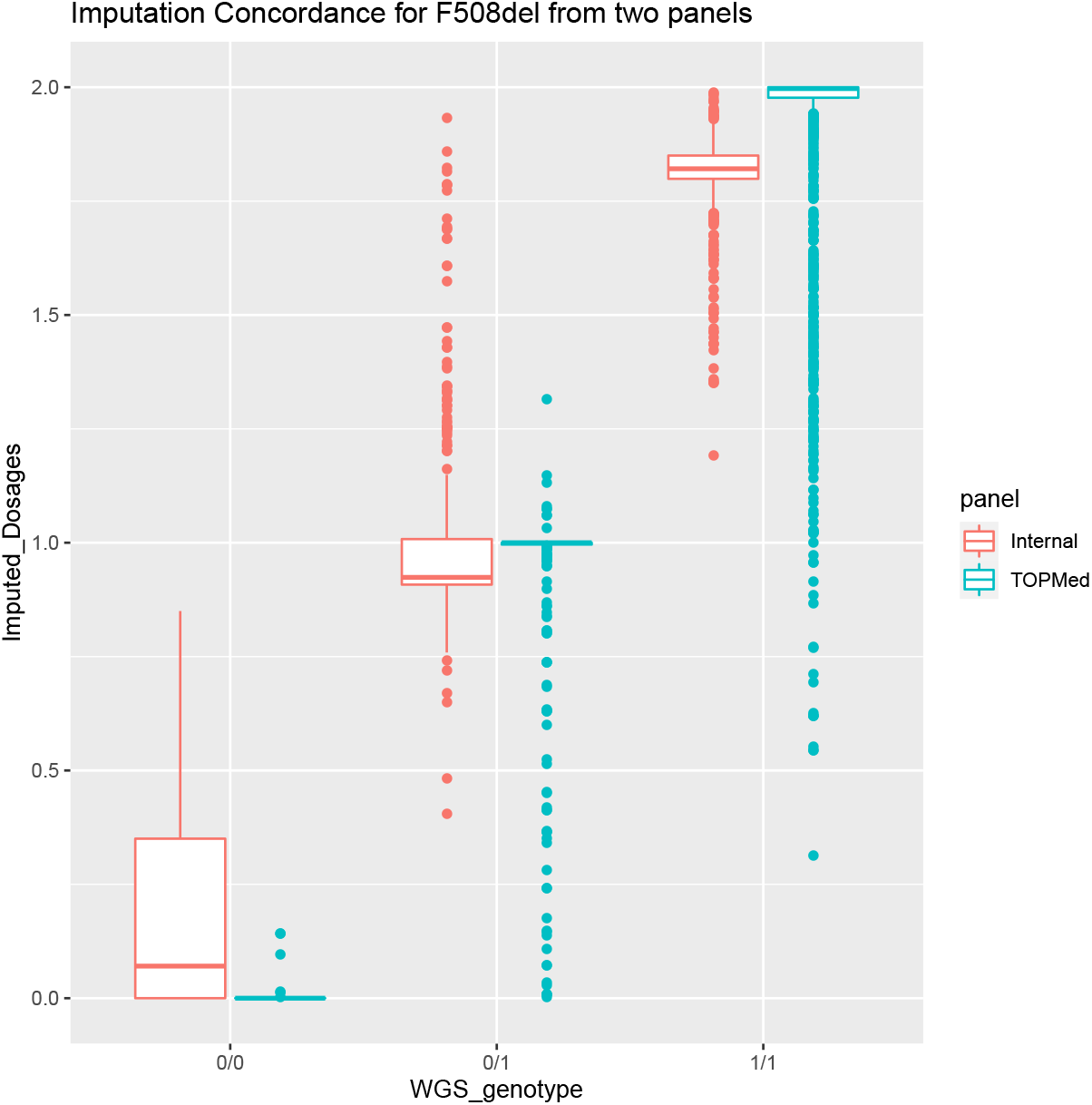
Imputation concordance for F508del using TOPMed and reduced CFGP reference panels. The true R2 for TOPMed and reduced CFGP imputed results are 0.835 and 0.926, and the sum of squared error for TOPMed and reduced CFGP are 117.58 and 82.42, respectively. The main reason that TOPMed is slightly worse is that it tends to under-estimate the deletion frequency.

Comparing with other imputation reference panels, we found the TOPMed reference panel provides much enhanced genome coverage. For example, for 610-Quad and 660W-set1 panels, TOPMed resulted in a 2.1-3.0x increase (**Table S2**) in genome coverage for LFRV compared with previous imputation using the Haplotype Reference Consortium (HRC) reference panel^7^. Overall, TOPMed-based imputation in CF patients is of satisfying quality, suggesting the value of TOPMed imputation reference panel for CF patients.

### Evidences showing value of constructing CFGP reference panel

Although publicly available genotype imputation reference panels from general populations (e.g. TOPMed freeze 8 reference panel) perform reasonably well for CF patients, we hypothesize that we may attain even better imputation quality for *CFTR* or other CF-associated loci by leveraging haplotype and linkage disequilibrium information among CF patients given rather drastic allele and haplotype differences in these regions between CF patients and general populations.

We performed Fisher’s exact test for each overlapped variant between CF WGS and TOPMed to compare the allele frequency difference between CF patients and general populations of >13,000 TOPMed participants of European ancestry from the TOP-LD project^19^, since over 95% of our CF patients are primarily of European ancestry. We found that *CFTR* gene and the region nearby is significantly enriched (p-value < 2.2e-16, **Table S3**) with variants with differential allele frequency (defined by Fisher’s exact test p-value < 2.5e-8 after Bonferroni correction) compared to other variants on chromosome 7. Previous work has also shown the benefit of cohort-specific reference panels^11,12^, including a study specifically targeted for CF patients^7^. With our WGS data with >5,000 samples, it is highly warranted to re-evaluate the utility of a CF-specific reference panel. To save some samples with WGS data for imputation quality evaluation, we constructed a *reduced CFGP reference panel* built from WGS data of 2,850 samples to impute another 1,992 unrelated samples to assess the value of a cohort-specific imputation reference panel.

### Imputation with reduced CFGP reference panel and quality evaluation

For the 1,992 samples, we compared their imputed data from the *reduced CFGP reference panel* (*n*=2,850) with that from the TOPMed freeze 8 reference panel (*n*=97,256). Note that TOPMed reference sample size is >34X that of the *reduced CFGP reference*. Not surprisingly, across all variants on chromosome 7 imputed by both reference panels, TOPMed clearly outperforms the *reduced CFGP reference panel* (**Figure 2A**), but the advantage becomes less pronounced when restricted only to the *CFTR* region (**Figure 2B**). Among the 544 *CFTR* variants, 138 are better imputed using the *reduced CFGP reference panel*, where 11/138 are highly damaging (CADD phred score^13^ > 20). This 8% (11/138) of highly damaging variants implies an 8X enrichment, because genome-wide we expect 1% of variants to be highly damaging based on the definition of CADD phred score where a score of 20 means among the 1% most damaging.

**Figure 2.**
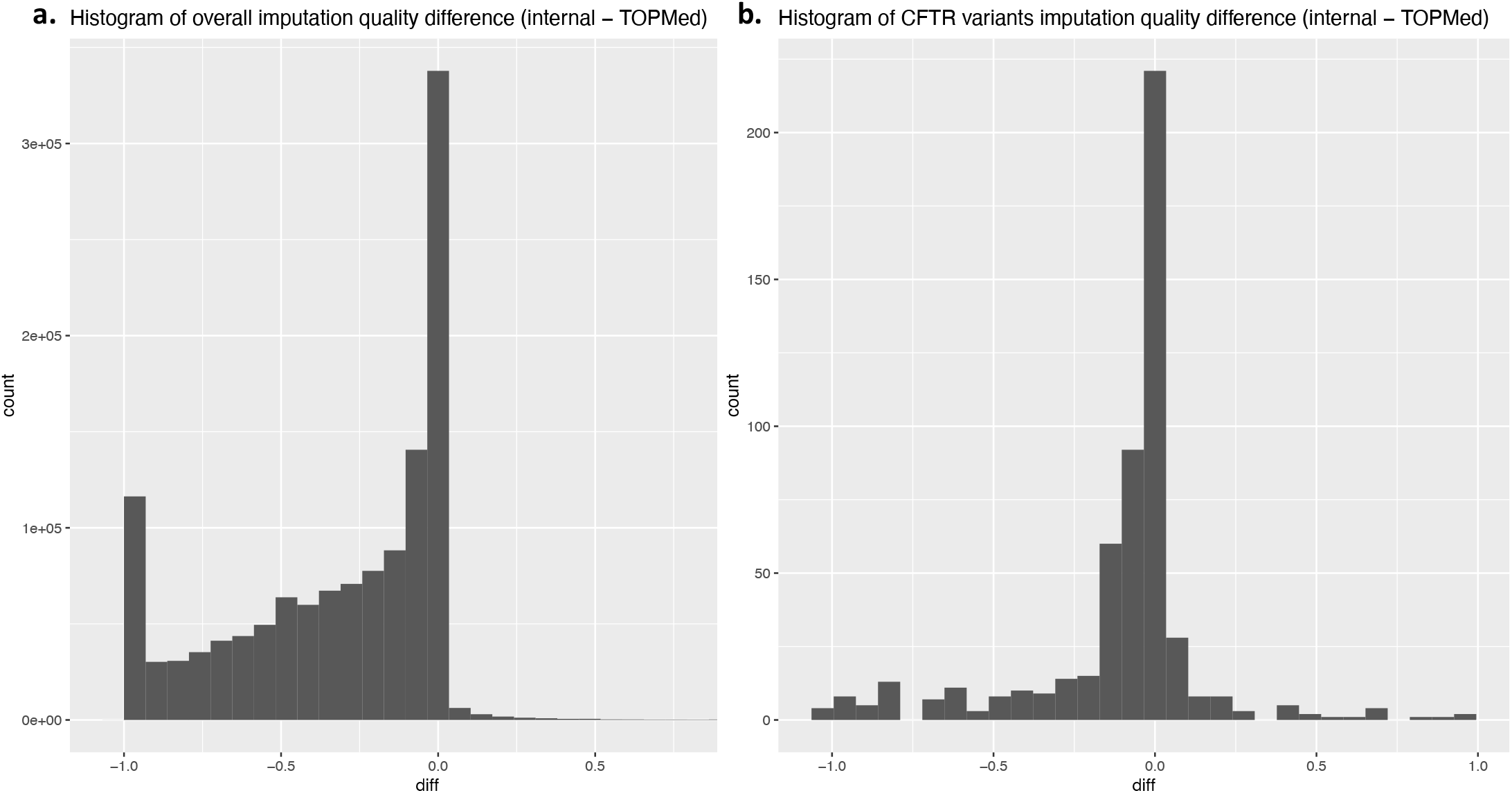
Histograms of differences between reduced CFGP true R2 and TOPMed true R2 to compare the imputation quality of the two reference panels. (A) For overall chr7. Almost all variants are located to the left half, which means TOPMed is predominantly better than the *reduced CFGP reference panel*. (B) For CFTR region only. The advantage of TOPMed reference panel over *reduced CFGP* becomes less pronounced.

Most of the *CFTR* variants that are much better imputed using the *reduced CFGP reference* panel are much rarer in TOPMed freeze 8 than among CF patients, explaining why the CF-specific reference panel leads to better performance. For example, for variant rs1244070394 (chr7:117480621:T:C, [GRCh38]), among the 132,345 TOPMed freeze 8 samples, we observe a MAC = 3 (MAF = 1.1e-5); while the MAC in our much smaller CFGP WGS samples (*n =* 5,095) is larger than that of TOPMed freeze 8: specifically MAC = 6, MAF = 5.9e-4. Although rare, some of these variants play important functional roles, with a few examples listed in **Table 4**. For instance, rs77284892 (chr7:117509047:G:T, [GRCh38], c.178G>A, p.Glu60Lys; legacy name E60K), with a MAF = 2.1e-3 in CFGP and MAF = 1.1e-5 in TOPMed freeze 8, has a CADD phred score of 38 (meaning the variant is among the 0.016% most deleterious variants in the human genome), is a stop-gain variant, and is classified as a CF-causing variant according to CFTR2. For the *CFTR* F508del variant, although the *reduced CFGP* imputation shows slightly larger bias than TOPMed imputation, it has a shorter tail and smaller variance, and is more consistent with true genotypes (**Figure 1**). The squared Pearson correlation between WGS true genotypes and *reduced CFGP* imputed dosages is 0.93, while that for TOPMed imputed dosages is 0.83. The long tail distribution of TOPMed imputed dosages for 1/1 homozygotes (i.e., homozygote deletion genotype) impedes its performance.

**Table 4.** Examples of variants that are much better imputed with reduced CFGP.

|  |  |  |  |  |  |
| --- | --- | --- | --- | --- | --- |
| **Variant (hg38)** | **chr7:1174806**<br>**21:T:C** | **chr7:1175090**<br>**47:G:T<sup>a</sup>** | **chr7:1175594**<br>**71:T:C<sup>a</sup>** | **chr7:1175877**<br>**38:G:A<sup>a</sup>** | **chr7:1176561**<br>**13:C:T** |
| rsIDs | rs1244070394 | rs77284892 | rs139573311 | rs76713772 | rs893051013 |
| CFGP true $R^2$ | 0.9934 | 0.9968 | 0.9703 | 0.9837 | 0.9423 |
| TOPMed true $R^2$ | 0.5490 | 0.3333 | 2.52e-7 | 0.7799 | 0.5010 |
| CF5095 AC | 6 | 21 | 8 | 115 | 21 |
| CF5095 AF | 5.89e-4 | 2.06e-3 | 7.85e-4 | 0.0113 | 2.06e-3 |
| TOPMed8 AC | 3 | 3 | 2 | 20 | 6 |
| TOPMed8 AF | 1.13e-5 | 1.13e-5 | 7.56e-6 | 7.56e-5 | 2.27e-5 |
| CADD phred score | 0.809 | 38 | 25.8 | 29.1 | 1.097 |
| VEP annotation | intron | stop gain | missense | splice acceptor | intron |
| CF-disease causing<sup>b</sup> | No | Yes | Yes | Yes | No |
| CFTR mutation | c.53+474T>C | c.178G>A<br>p.Glu60Lys | c.1400T>C<br>p.Leu467Pro | c.1585-1G>A | c.3963+3182C<br>>T |

Abbreviations are as follows: AC, allele count; AF, allele frequency.
<sup>a</sup> The middle three variants have very high CADD phred scores and are disease causing variants, but their TOPMed imputation qualities are not satisfying. It shows the value of our CF-specific reference panel.
<sup>b</sup> According to cftr2.org

We also broke down these variants by functional categories (simply coding and non-coding) to see whether the *reduced CFGP* reference panel performs better for functionally important variants. Due to the small number of coding variants, we didn’t further split the coding category. As expected, the *reduced CFGP* reference panel performs better for coding variants than non-coding variants, but less well compared to TOPMed (**Table S5**). However, the *χ*^2^ test shows variants that were better imputed with *reduced CFGP* is significantly enriched with coding variants (p = 5.5e-3, OR = 2.61). We also found the reduced CFGP reference panel performs better for less common variants compared to common variants, but TOPMed still outperforms the reduced CFGP for the vast majority due to the large sample size difference (Table S6).

We then systematically compared the performances of the two reference panels across the whole genome to see whether the *reduced CFGP* reference panel performs better in any genome regions other than the *CFTR* region on chromosome 7. Specifically, we calculated the difference of *reduced CFGP* imputed true R2 and TOPMed imputed true R2 (the former minus the latter) for each variant, and then summarized variant level true R2 difference at 1MB non-overlapping region level. We used two statistics for region-level summary: mean true R2 difference of variants 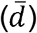 and the proportion of variants whose true R2 difference is greater than 0 (*p*) indicating that *reduced CFGP* performs better than TOPMed, in the corresponding 1MB region. To increase stability, we only considered regions harboring over 100 variants for evaluations. For the whole genome, 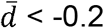 and *p* < 8% for most of the 1MB regions (**Figure 3**). As a positive control, for the *CFTR* region, 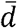 ranges from -0.2 to -0.13, and *p* ranges from 12% to 20%, with each statistic falling in the 1% of its distribution. Interestingly, some other regions show even stronger evidence that the relative (to TOPMed) performance of the *reduced CFGP* reference panel is substantially better than genome-average, including the 60-66 MB region on chromosome 9 (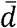 ranges from -0.17 to -0.09, *p* ranges from 28% to 33%), 19-23 MB region on chromosome 15 (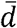 ranges from -0.06 to -0.03, *p* ranges from 21% to 29%), as well as the HLA region (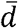 ranges from -0.15 to -0.10, *p* ranges from 11% to 18%) (**Table S7**). We currently do not fully understand why the relative performance of *reduced-CFGP* reference panel over TOPMed in these regions are better than genome-average. The regions do not seem to colocalize with known GWAS loci because these outlier regions we identified are not close to reported GWAS signals and regions harboring known GWAS variants do not show large 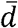 or *p* compared to genome-average. The region-level summary statistics are tabulated in **Table S7** for other researchers to further investigate.

**Figure 3.**
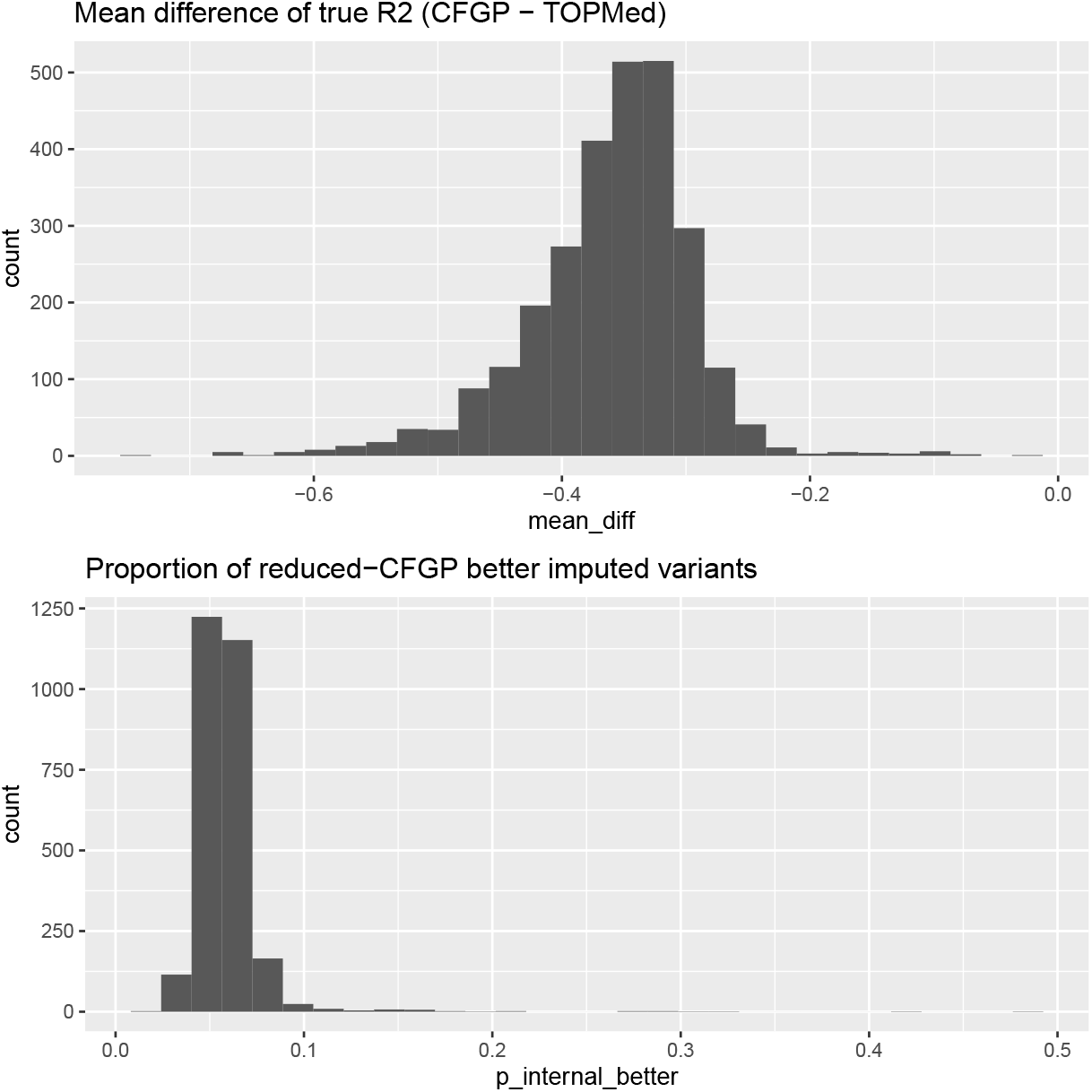
Histograms of mean true R2 difference and proportion of variants better imputed by reduced CFGP than TOPMed, across 2872 1Mb non-overlapping regions. We calculated the true R2 difference of the two reference panels using reduced-CFGP true R2 minus TOPMed true R2 for each variant, and then summarized variant level true R2 difference at 1Mb region level using the two statistics.

This proof-of-concept experiment showcases the value of a CF-specific reference panel for imputing data for CF patients, particularly in some specific regions (e.g. the *CFTR* region), on top of the state-of-the-art TOPMed reference panel. Thus, we constructed a *CFGP reference panel* using the full set of 5,095 WGS samples in the CFGP. We anticipate this *CFGP reference panel* to be valuable for other investigators studying CF but having only array density genotype data instead of WGS data.

### Imputation improves PRS performance

We further constructed polygenic risk scores (PRS) for KNoRMA^14^ to assess whether imputation, particularly TOPMed-based imputation, would help construct a PRS with higher prediction accuracy. KNoRMA is a quantitative lung trait of FEV1 data over 3 years adjusted for survival^14^ measuring lung function, and is one of the main focused traits in the CFGP consortium. PRS are usually constructed as weighted summation of genetic markers, where the weights are derived from GWAS in independent training samples. Here, we hypothesize that imputation would improve PRS performance, either by imputing target samples where PRS formula is applied to, or by imputing training samples where GWAS is performed to construct the PRS formula. We performed two experiments to mimic two realistic scenarios: (1) whether imputation is performed in the target cohorts where PRS is applied to (**Figure 4A**); (2) whether imputation is performed in the discovery cohorts where the PRS is constructed (**Figure 4B**). In the second scenario, we have some samples WGSed and others only genotyped with some genotyping array to start with. We then compared the accuracy of PRS constructed with or without imputation.

**Figure 4.**
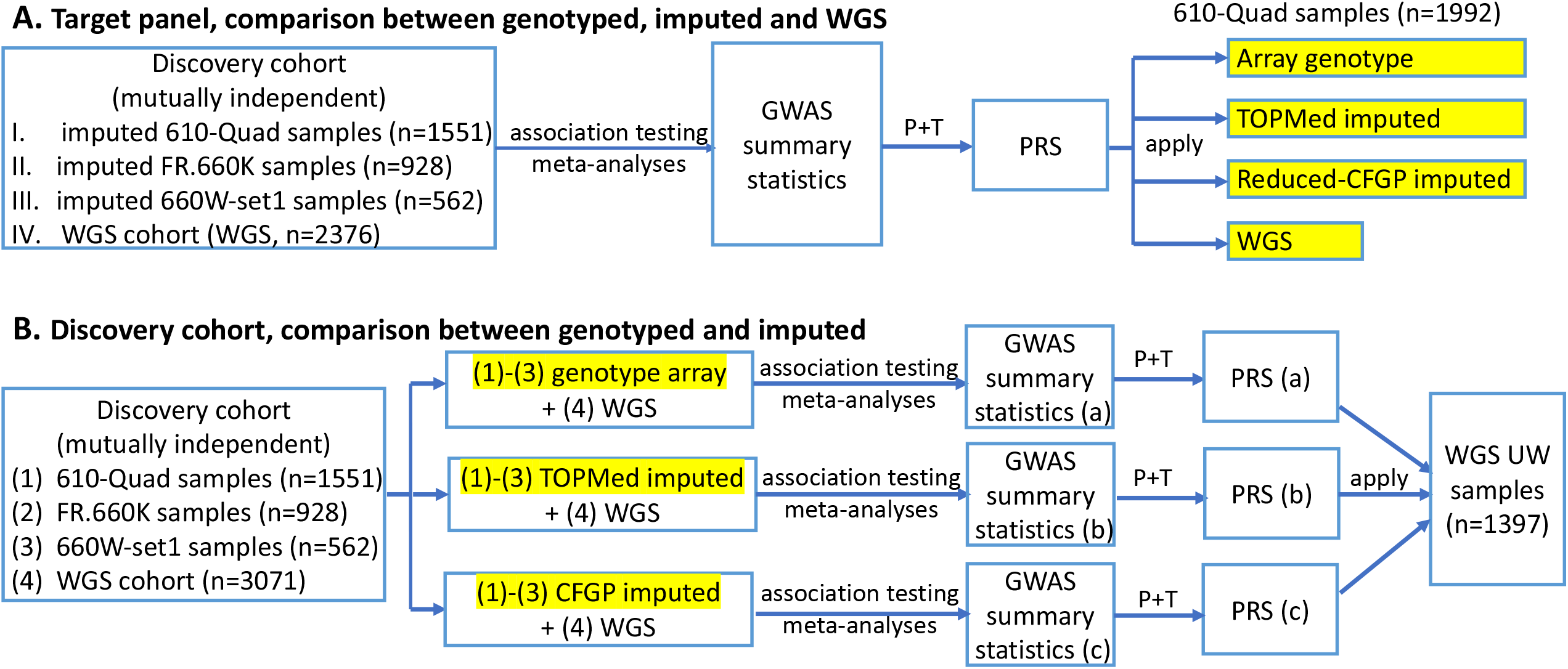
Illustration of impact of imputation on PRS construction. **A. Imputation performed in target cohorts.** We started with four independent discovery cohorts (I-III are TOPMed imputed data, IV is WGS data), performed association analysis for each subset separately and then meta-analyzed the association results. The meta-GWAS summary statistics was then used to construct PRS using the P+T method. The constructed PRS was applied to the same 1992 target samples but with four different marker densities (in yellow highlight): array genotype, TOPMed imputed, Reduced-CFGP imputed or WGS data to compare the benefit of imputation in target cohort. **B. Imputation performed in discovery cohorts**. We started with the same first three discovery cohorts as in A but adopted three different marker sets (again in yellow highlight), as well as a fourth independent WGS cohort. We then performed association analysis and meta-analysis for each marker set, and constructed three different PRSs using the three different meta-GWAS summary statistics. The three PRSs were then applied to the same cohort to compare the performances.

To test the benefit of imputation for PRS target cohorts, we applied the same PRS to the 1992 samples for whom we have 610-Quad array, TOPMed-based imputation and *reduced CFGP* based imputation (both starting from 610-Quad array), and WGS data available. The PRS was constructed based on GWAS summary statistics from meta-analysis of samples independent of the 1992 test samples (**Figure 4A, Methods Section A**). Four different marker sets (genotype array data only, TOPMed imputed data with Rsq > 0.3, *reduced CFGP* imputed data with Rsq > 0.3 and WGS data) were adopted for the application of PRS. We performed a grid search over MAF and p-value threshold (**Methods**) and reported the best one (largest correlation with true KNoRMA values after adjusting for age, sex, study, and first 6 PCs) to compare the four different marker sets. We found that with TOPMed imputation, we can nearly achieve the same performance as WGS (**Table S4**). The PRS correlation improves by 37.2% with TOPMed imputation compared to genotype array data only, while only 0.99% inferior to WGS data. The *reduced CFGP* imputed data also performs satisfactorily, especially considering the much smaller reference panel size. It improves the PRS correlation by 32.1% compared to genotype array data only, while only 4.7% inferior to WGS data.

To evaluate the benefit of imputation in PRS discovery/construction cohorts, we took UW samples (n=1397) with only WGS data as the target cohort, and applied three different sets of PRSs (**Figure 4B**). The three different sets of PRSs differ by the marker density in the same discovery cohorts consisting of 6,112 samples independent of the UW samples (**Figure 4B, Methods Section B**). Specifically, the first set of PRS was constructed based on association summary statistics from meta-analyzing 3,041 patients with array data and 3,071 patients with WGS data (**Figure 4B (a)**). The second and the third sets were constructed similarly, only replacing the 3,041 patients from array data to TOPMed-imputed (**Figure 4B (b)**) or CFGP-imputed data (**Figure 4B (c)**). We similarly compared the best PRS searched over different MAF and p-value threshold grids under the three different sets of GWAS summary statistics, finding the TOPMed-imputation-aided PRS results in 71.2% higher correlation, while the CFGP-imputation-aided PRS results in only 9.0% higher correlation, compared to that without imputation (**Table 5**). We further performed two-sample t-test to compare the KNoRMA values of samples from top and bottom 5% of predicted PRS, to test the power of the three PRS sets in stratifying patients in terms of lung function gauged by KNoRMA values. We found significant difference in KNoRMA value for patients from two extreme tails predicted by the imputation-aided PRS (p-value = 0.038 for TOPMed-based imputation and p-value = 0.0065 for CFGP-based imputation), while no significant difference in the PRS without imputation counterpart (p-value = 0.712) (**Table 5**).

**Table 5.**
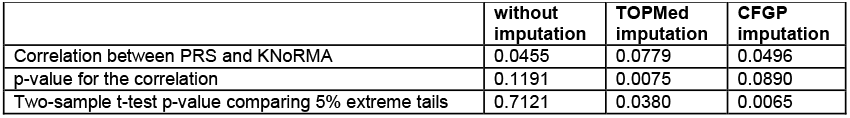
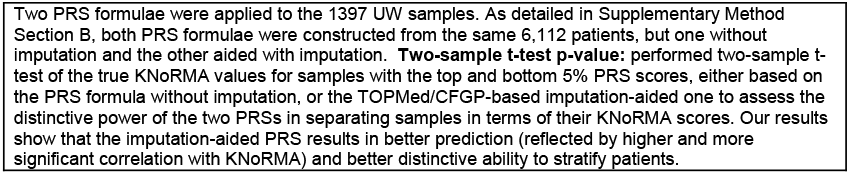
PRS performance when applied to UW samples.

|  | without imputation | TOPMed imputation | CFGP imputation |
| --- | --- | --- | --- |
| Correlation between PRS and KNoRMA | 0.0455 | 0.0779 | 0.0496 |
| p-value for the correlation | 0.1191 | 0.0075 | 0.0890 |
| Two-sample t-test p-value comparing 5% extreme tails | 0.7121 | 0.0380 | 0.0065 |

Two PRS formulae were applied to the 1397 UW samples. As detailed in Supplementary Method Section B, both PRS formulae were constructed from the same 6,112 patients, but one without imputation and the other aided with imputation. **Two-sample t-test p-value:** performed two-sample t-test of the true KNoRMA values for samples with the top and bottom 5% PRS scores, either based on the PRS formula without imputation, or the TOPMed/CFGP-based imputation-aided one to assess the distinctive power of the two PRSs in separating samples in terms of their KNoRMA scores. Our results show that the imputation-aided PRS results in better prediction (reflected by higher and more significant correlation with KNoRMA) and better distinctive ability to stratify patients.

## Discussions

In summary, even for patients affected with a Mendelian disease as CF, TOPMed reference panel leads to satisfactory genome-wide imputation quality, and better PRS prediction accuracy. We further demonstrate the value of a CF-specific reference panel, which can outperform TOPMed for some variants due to better match with target (also CF) samples in terms of allele and haplotype frequencies. Although at 1Mb region level, a CF-specific reference panel never outperformed TOPMed reference panel, in some regions, it offers substantially more complementary information to TOPMed. These regions include the *CFTR* region harboring the gene causing this Mendelian diseases, and several other genome regions including HLA. Our CFGP reference panel consisting of >10,000 haplotypes developed from WGS data from CF patients should benefit other investigators in their genetic studies of CF.

We note that the value demonstrated in our experiments with *reduced CFGP* reference panel is not simply due to samples from the same recruitment sites between references and targets. The 1,992 samples as targets were from three different studies (CGS, GMS, TSS), and the 2,850 samples as reference were from four different studies, including an independent study, EPIC, in addition to the three studies. In order to show that the performance of disease-specific CF panel is not due to overlapping of samples from the same recruitment sites, we additionally performed imputation for the same 1,992 target samples using EPIC-only samples as reference. In this case, samples in targets and references are from completely independent recruitment sites. We then plotted the histograms of imputation quality difference between different reference panels and found most of the variants exhibit highly similar qualities and the EPIC-only reference panel similarly leads to a larger proportion of variants around *CFTR* better imputed than when using TOPMed as the reference (**Figure S2 c**,**d**). These results demonstrate that the benefit is not simply due to overlapping of samples from the same recruitment sites, but the similarity of genomes in CF patients. Furthermore, our study would not only benefit the CF community, but also provide a genotype imputation protocol for other Mendelian diseases. With more WGS data in production, future investigators studying other Mendelian diseases could further explore benefits of disease-specific imputation reference panels.

Since cohort-specific reference panel provides better match in terms of allele and haplotype frequencies, while TOPMed reference panel benefits from its much larger sample size, future work can further explore strategies to combine the two reference panels. Directly combining different reference panels is largely infeasible due to different marker densities and restricted access to individual-level haplotypes. An alternative approach is to combine two or more sets of imputed results using “meta-imputation”, which outputs a consensus imputed dataset by calculating weighted sum of single-reference imputed results, such as implemented in MetaMinimac2. Another direction is to perform marker-level selection of reference panels, where the issue is that we cannot easily quantify the relative performance of reference panels without true genotypes. In our study, we found the state-of-the-art imputation quality estimation metric, Rsq output by minimac, tends to favor the TOPMed reference panel, even when the true quality from *reduced CFGP* reference panel is much better than that from TOPMed. For example, for the last variant in **Table 4**, rs893051013 (chr7:117656113:C:T, [GRCh38]), selection of reference panel based on Rsq would strongly favor TOPMed (Rsq is 0.80, much higher than 0.29 from the *reduced CFGP*), but in reality the *reduced CFGP* performed much better: true R2 achieved 0.94, much better than TOPMed resulting in a true R2 of only 0.5. Future research should explore imputation quality metric that either more accurately reflect true quality or at least comparable across reference panels.

Besides providing further enhanced imputation reference panels, WGS is also valuable in many other aspects, including enabling the study of variants other SNPs and more comprehensively identifying disease causing variants. As one example, for the 281 disease causing variants reported by CFTR2 that can be mapped to GRCh38 positions, CFGP WGS data covered 137 of them, while only 35 were well-imputed by TOPMed, demonstrating the value of generating WGS data for the CF community. Although 25.5% (35/137) is not ideal, imputation substantially enhances over genotyping array with 1-10 of these 137 variants directly genotyped, or over earlier imputation references panels (e.g., with 1000 Genomes reference, 15 out of the 137 variants can be well imputed). Therefore, before WGS data is available for every CF patient, imputation using TOPMed or CFGP reference panel provides a substantial boost.

## Methods

### Genotype array data and pre-imputation quality control (QC)

There are in total 7,988 samples genotyped on seven different arrays before QC (**Table S1**). Note that there are some duplicates/triplicates, thus the 7,988 samples represent < 7,988 unique patients. We will not get into the patient level in this paper. since one patient can contribute to more than one samples, either through recruitment by more than one study site, or by being genotyped more than once. All the imputation metrics reported were calculated at sample level.

We performed both sample-and variant-level QC prior to imputation. We removed samples with genotype missing rate > 10% using plink v.1.90. 18 samples in the arrays were excluded due to this low call rate criterion. We further removed unexpected alleles (e.g., N), monomorphic sites, ambiguous SNPs (A/T or C/G SNPs) and then lifted over from hg19 to hg38. The final numbers of QC+ variants in each GWAS array ranged from 263,660 to 3,379,381 (**Table S1**).

### TOPMed imputation

We first performed strand flipping according to our reference panel (TOPMed Freeze 8) to improve imputation accuracy. Ambiguous SNPs (i.e., A/T or C/G SNPs) had already been dropped in the pre-imputation QC step above. For non-ambiguous SNPs, the alleles in our cohort were flipped if they appear in minus strand, when compared to the reference panel (for example, the alleles in our cohort are A/G, while they are T/C or C/T in the reference panel). We used the TOPMed Imputation Server (https://imputation.biodatacatalyst.nhlbi.nih.gov/#!) for phasing (via eagle^15^) and imputation (via minimac4^16^), using the TOPMed freeze 8 as the reference panel. This reference panel, built from 97,256 deeply sequenced human genomes, contains 308,107,085 genetic variants. After imputation, we retained only variants with imputation quality (Rsq or estimated R2) ≥ 0.3.

### True imputation quality metric (trueR2)

We calculated the true imputation quality metric (true R2, the squared Pearson correlation between imputed dosages and true genotypes with the latter coded as 0, 1 and 2) to evaluate our imputation quality. The true genotypes were derived from the CFGP WGS data. We first intersected our imputed variants with WGS PASS variants by MAF bins (here, “true” MAF as defined by genotypes derived from WGS data). Then, we extracted the genotypes for overlapped samples between GWAS and WGS to evaluate the concordance. Our evaluation was restricted only to samples with QC+ data from GWAS and WGS. Duplicate samples were also dropped. Finally, the squared Pearson correlation was calculated for each variant, which is the true R2. Note that this true R2 is different from estimated R2 or Rsq above in that estimated R2 or Rsq is part of the imputation output and is obtained in the absence of true genotypes. By contrast, true R2 can only be calculated when the true genotypes are available, which is not realistic except for evaluation purposes because if we had true genotypes, we would not have bothered with imputation.

### Imputation based on a *Reduced CFGP Reference Panel*

As a proof-of-concept experiment, we constructed a *reduced CFGP* imputation reference panel using WGS data of 2,850 samples from the CF Genome Project (CFGP). Such reference construction has been commonly adopted, particularly when target samples (i.e., samples to be imputed) do not match well with those in standard imputation reference panels. We started with QC+ WGS data and performed phasing using eagle^15^ with default parameters to generate the *reduced CFGP reference panel*. Using our self-constructed *reduced CFGP reference panel*, we imputed chromosome 7, where *CFTR*, the CF causing gene, is located, in 1,992 samples, independent of the 2,850 samples contributing the *reduced CFGP reference panel*. These 1,992 samples have WGS data and have also previously been genotyped on the 610-Quad array with 30,853 QC+ GWAS markers on chromosome 7. We assessed the relatedness between this target sample of 1,992 samples and the 2,850 samples in the *reduce CFGP reference panel* using plink --genome. Distribution of the PI_HAT is shown in (**Figure S1**) with the maximum PI_HAT < 0.1. With the low level of relatedness between target and reference, we proceeded with imputation in the target sample using minimac4^16^ with default parameters and compared the imputed dosages with true genotypes derived from their WGS data.

To evaluate the value of the *CFGP reference panel* in comparison to commonly used imputation reference panels, we also compared the performance of the *CFGP reference panel* relative to the state-of-the-art TOPMed freeze8 reference panel.

### Construction of a CFGP reference panel

Similar to the *reduced CFGP* reference panel, the *CFGP reference panel* was constructed from CFGP WGS data. Different from the *reduced CFGP reference* where a subset of 2,850 samples were used, the *CFGP reference* was built from all 5,095 samples in CFGP. We similarly started with QC+ WGS and constructed the CFGP reference by phasing with eagle with default parameters.

### Generating genome-wide association statistics for PRS construction

GWAS were performed separately for different subsets of samples using the EMMAX test implemented in EPACTS v3.3.0^17^, which accounts for genetic relatedness via a mixed model approach. Specifically, the model adjusts for a kinship matrix that was calculated using genotyped variants with missing rate < 1% and MAF > 1%. When performing the association testing, we restricted to variants with MAF > 0.1% and imputation Rsq > 0.3 when running EPACTS to improve model stability. In each subset GWAS analysis, we adjusted for age, sex, study and first 6 PCs. We then used METAL^18^ for meta-analysis to enhance the discovery sample size for improved power.

We note that the PRS construction seems complicated. The primary reason is the complicated data structure we have (several different genotype array datasets, and the mixture of array data, imputed data with two different reference panels, and WGS data). The idea in the section is rather straightforward: since PRS construction involves both training samples (where GWAS are performed and weights for PRS are derived) and independent target samples (where PRS formula is applied to and evaluated), we hypothesize that imputation in either target samples (**Figure 4A**) or training samples (**Figure 4B**) would improve the PRS performance in target samples. Figure 4A is the scenario where the only difference is the genetics data of target samples used when applying the PRS formula. We used array-only genotypes, TOPMed imputed data, CFGP imputed data and or WGS data in target samples, and evaluated the PRS calculated with the four different types of genetics data. Figure 4B is the scenario where the only difference is the genetics data of (part of the) training samples used when performing GWAS and to derive variant-specific weights forconstructing the PRS formula. We used array array-only genotypes, TOPMed imputed data, and or CFGP imputed data in (part of the) training samples when deriving the PRS weights. We say “part of the” training samples because for all three settings in **Figure 4B**, we used WGS for the 3,071 samples with WGS data.

**Section A**. For experiments where the 1992 610-Quad samples with both array and WGS data are used as target samples, the discovery cohorts include the following four sets of 5,417 samples, all independent of the target 1992 samples: (1) 610-Quad samples (n=1551, TOPMed imputed); (2) FR.660K samples (n=928, TOPMed imputed); (3) 660W-set1 samples (n=562, TOPMed imputed); and (4) WGS samples (n=2376, WGS data).

**Section B**. For experiments where the 1397 UW samples with WGS data are used as target, the discovery cohorts include the following four sets of sample, similarly all independent of the target 1397 UW samples (1) 610-Quad samples (n=1551, genotyped or TOPMed/CFGP imputed); (2) FR.660K samples (n=928, genotyped or TOPMed/CFGP imputed); and (3) 660W-set1 samples (n=562, genotyped or TOPMed/CFGP imputed); and (4) WGS samples other than UW (n=3071, WGS data). The summary statistics without imputation refers to (1)-(3) with array genotype + (4) when conducting associations (**Figure 3B (a)**),, the summary statistics with TOPMed imputation refers to (1)-(3) with TOPMed imputed data + (4) when conducting associations (**Figure 3B (b)**), and the summary statistics with CFGP imputed refers (1)-(3) with CFGP imputed data + (4) when conducting associations (**Figure 3B (c)**).

### PRS construction

We constructed PRS with the common P+T method performed with plink v1.90. We performed a grid-search over different MAF (≥0.1%, ≥0.5%, ≥1%, ≥5%) and p-value thresholds (≤1, ≤0.5, ≤0.1, ≤0.05, ≤0.01, ≤5e-3, ≤1e-3, ≤5e-4, ≤1e-4, ≤5e-5, ≤1e-5) combinations to determine the best performance under each different target or discovery marker sets. For chromosome X, males were coded as 0 or 2.

## Web resources

1. TOPMed imputation server: https://imputation.biodatacatalyst.nhlbi.nih.gov/#!
2. Eagle: https://alkesgroup.broadinstitute.org/Eagle/
3. Minimac4: https://genome.sph.umich.edu/wiki/Minimac4
4. Bravo: https://bravo.sph.umich.edu/freeze8/hg38/
5. CFTR2: https://cftr2.org
6. plink v1.90: https://www.cog-genomics.org/plink/1.9/
7. EPACTS: https://genome.sph.umich.edu/wiki/EPACTS
8. TOP-LD: http://topld.genetics.unc.edu/topld/index.php
9. MetaMinimac2: https://github.com/yukt/MetaMinimac2

## Acknowledgement

This work is supported by CFF grants CUTTIN18XX1, BAMSHA18XX0, KNOWLE18XX0 and KNOWLE21XX0, and is submitted on behalf of the CF Genome Project. Additional support from NHLBI BioData Catalyst Fellowship awarded to Jia Wen: 1OT3HL142479-01, 1OT3HL142478-01, 1OT3HL142481-01, 1OT3HL142480-01, 1OT3HL147154.

We would like to thank the Cystic Fibrosis Foundation for the use of CF Foundation Patient Registry data to conduct this study. Additionally, we would like to thank the patients, care providers, and clinic coordinators at CF centers throughout the United States for their contributions to the CF Foundation Patient Registry.

Furthermore, we would like to acknowledge use of the Trans-Omics in Precision Medicine (TOPMed) program imputation panel (freeze 8 version) supported by the National Heart, Lung and Blood Institute (NHLBI); see www.nhlbiwgs.org. TOPMed study investigators contributed data to the reference panel, which was accessed through https://imputation.biodatacatalyst.nhlbi.nih.gov. The panel was constructed and implemented by the TOPMed Informatics Research Center at the University of Michigan (3R01HL-117626-02S1; contract HHSN268201800002I). The TOPMed Data Coordinating Center (R01HL-120393; U01HL-120393; contract HHSN268201800001I) provided additional data management, sample identity checks, and overall program coordination and support. We gratefully acknowledge the studies and participants who provided biological samples and data for TOPMed.

## Data Availability

The CFGP WGS data are available for request to the Cystic Fibrosis Foundation at https://www.cff.org/researchers/whole-genome-sequencing-project-data-requests#requesting-data.

