## Supplementary Materials for "Leveraging TOPMed Imputation Server and Constructing a Cohort-Specific Imputation Reference Panel to Enhance Genotype Imputation among Cystic Fibrosis Patients"

**Supplementary Tables and Figures**

| Illumina Panel | # samples | # variants | # samples QC+ | # variants QC+ |
| --- | --- | --- | --- | --- |
| FR.300K | 144 | 263,660* | 144 | 263,660 |
| FR.370K | 145 | 309,012 | 145 | 308,937 |
| FR.660K | 1,011 | 554,657 | 1,011 | 552,744 |
| 610-Quad | 3,844 | 570,663 | 3,840 | 567,784 |
| 660W-set1 | 2,026 | 655,214 | 2,012 | 556,532 |
| 660W-set2 | 444 | 655,214 | 444 | 551,819 |
| Omni5 | 374 | 4,289,087 | 374 | 3,379,381 |

**Table S1. Summary of seven GWAS arrays.**

*FR.300K was originally combined into FR.370K. We did not realize that until lifting over. The number shown here is after removing unexpected alleles, monomorphic sites, non-biallelic variants, and further lifting over to hg38. The procedure is demonstrated in the Supplementary Method.

| Reference panel and GWAS array | # Total variants | # variants with Rsq >= 0.3 | # variants with MAF < 0.5% | # variants with Rsq>=0.3 and MAF < 0.5% | fold increase comparing to HRC |
| --- | --- | --- | --- | --- | --- |
| HRC previous work | 2,283,806 | 1,375,928 | 1,754,169 | 850,266 | - |
| TOPMed 8 601-Quad | 16,990,285 | 3,167,307 | 16,376,277 | 2,561,141 | 3.0 |
| TOPMed 8 660W-set1 | 16,990,285 | 2,435,712 | 16,367,583 | 1,819,447 | 2.1 |

**Table S2. Comparing TOPMed imputation genome coverage with previous reports for chromosome 7.** TOPMed freeze 8 reference panel can achieve 2.1-3.0 fold increase for well-imputed low frequency or rare variants.

|  | # variants with differential AF | # variants in total | Odds ratio (95% CI) | p-value |
| --- | --- | --- | --- | --- |
| CFTR variants | 354 | 827 | 4.14 (3.65, 4.70) | <2.2e-16 |
| 20MB bin CFTR locates | 12,251 | 231,466 | 2.12 (2.08, 2.16) | <2.2e-16 |
| All chr7 | 55,957 | 2,239,582 |  |  |

**Table S3. Assessment of variants with differential allele frequency in CF patients and TOPMed European ancestry samples.** We performed Fisher’s exact test for each variant overlapped between CF WGS and TOPMed, resulting in ~2.2 million variants in total. We define variants with differential AF as the p-value of Fisher’s exact test is less than 2.5e-8 after Bonferroni correction. CFTR variants refer to variants with position between 117,480,025 and 117,668,665. We further partitioned chr7 into 8 disjoint continuous 20MB bins to compare the enrichment of variants with differential AF variants in each bin. The 20MB bin with *CFTR* gene is 100-120MB. All positions are in hg38. We observe significant enrichment of variants with differential AF for *CFTR* gene and the 20MB bin it locates.

|  | array genotype only | TOPMed imputed  (Rsq > 0.3) | *Reduced-CFGP* imputed  (Rsq > 0.3) | WGS |
| --- | --- | --- | --- | --- |
| Correlation between PRS and KNoRMA | 0.0443 | 0.0608 | 0.0585 | 0.0614 |
| p-value for the correlation | 0.0555 | 0.0085 | 0.0114 | 0.0078 |

**Table S4. PRS performance when testing in 1992 610-Quad samples.** These 1992 samples have both genotype array (the Illumina 610-Quad array) and WGS data available. PRS was constructed from the 5,417 samples in Section A. We then applied the same PRS formula to four different sets of variants in our target 1992 samples: array genotype only, TOPMed imputed with Rsq > 0.3, *Reduced-CFGP* imputed with Rsq > 0.3 and WGS data. As expected, WGS performs the best, but TOPMed imputed sets nearly achieve the WGS performance. Three sets of PRS are significantly associated with true KNoRMA except the one when only genotype array data are used.

| Functional category | # total variants | # reduced CFGP better imputed variants | # TOPMed better imputed variants | % reduced CFGP better imputed variants |
| --- | --- | --- | --- | --- |
| non-coding | 504 | 120 | 384 | 23.8% |
| coding | 40 | 18 | 22 | 45% |

**Table S5. Imputation comparison of TOPMed and *reduced CFGP* reference panels for coding and non-coding variants.** The $\chi^{2}$ test shows variants that were better imputed with *reduced CFGP* is significantly enriched with coding variants (p = 5.5e-3, OR = 2.61)

| MAC range | # total variants | # reduced CFGP better imputed variants | # TOPMed better imputed variants | % reduced CFGP better imputed variants |
| --- | --- | --- | --- | --- |
| (0, 10] | 111 | 29 | 82 | 26.1% |
| (10, 20] | 77 | 34 | 43 | 44.2% |
| (20, 50] | 130 | 31 | 99 | 23.8% |
| (50, 200] | 56 | 18 | 38 | 32.1% |
| 200+ | 170 | 26 | 144 | 15.3% |

**Table S6. Imputation comparison of TOPMed and *reduced CFGP* reference panels breaking down by MAC range.** For less common variants, the *reduced CFGP* reference panel performs better compared to more common variants, but TOPMed still beats the *reduced CFGP* for the most majority due to the large sample size difference.

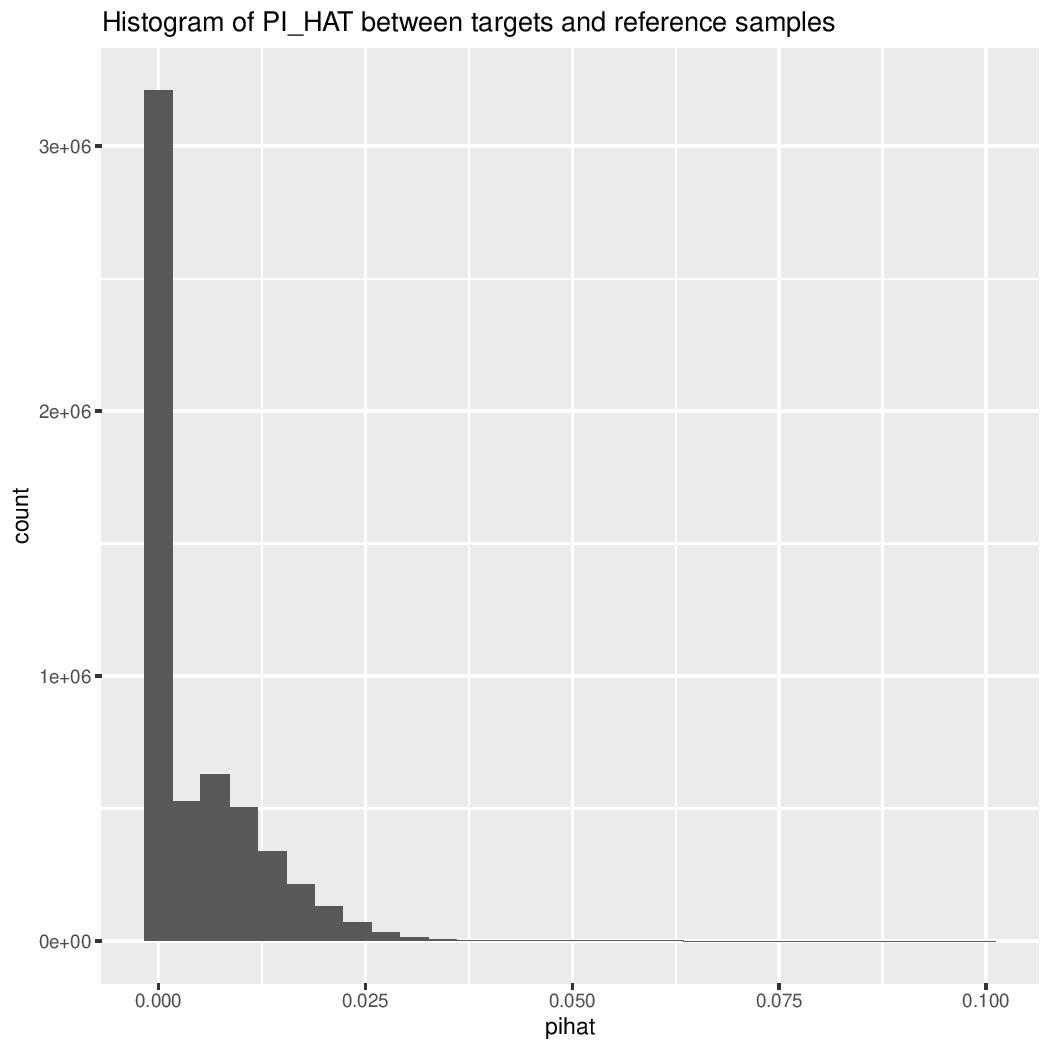

**Figure S1. The histogram of PI_HAT values of all the target-reference sample pairs.** Most sample pairs have no relationship at all (PI_HAT ~ 0), with the maximum value of 0.1. We confirmed that there are no relatedness issues to avoid over-estimate of the imputation quality using *reduced CFGP reference panel*.

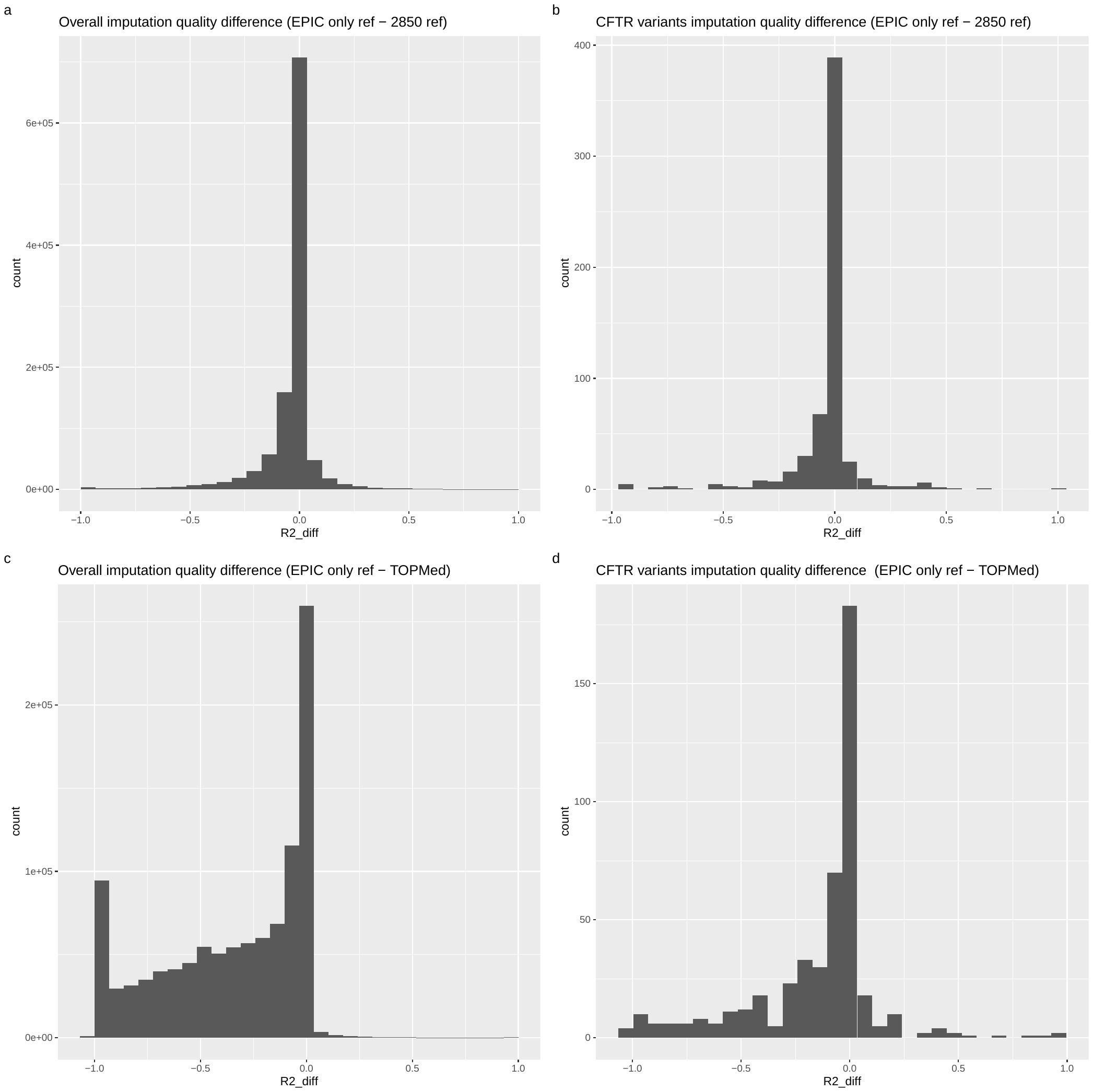

**Figure S2. The histogram of imputation quality between different reference panels. a & b**. Comparison between EPIC-only reference panel (n=1246) and reduced CFGP reference panel (n=2850) for the entire chromosome 7 (**a**) and the *CFTR* region only (**b**). The EPIC-only performs worse largely due to smaller reference size compared to the 2850 reference panel. **c & d**. Comparison between EPIC-only reference panel (n=1246) and TOPMed (n=97,256) for the entire chromosome 7 (**c**) and the *CFTR* region only (**d**). The EPIC-only reference panel is also comparable to TOPMed for the *CFTR* region, especially in contrast to the quality difference for the whole chromosome 7.
